# Proportion of cocaine-coding neurons in the orbitofrontal cortex determines individual drug preferences

**DOI:** 10.1101/050872

**Authors:** Karine Guillem, Serge H. Ahmed

**Author notes:** Correspondence should be addressed to K.G. or S.H.A.

## Abstract

Cocaine addiction is a harmful preference for drug use over and at the expense of other nondrug-related activities. Here we identify in the orbitofrontal cortex (OFC) – a prefrontal region involved in choice and decision-making – a mechanism that explains individual preferences in rats between cocaine use and an alternative, nondrug action. We found that initiation of these actions is selectively encoded by two non-overlapping populations of OFC neurons, and that the relative size and differential pre-choice activity of the cocaine action-coding population determine an individual preference, a larger size and a higher pre-choice activity being associated with cocaine preference. A larger size is a structural feature that may confer to a population of OFC neurons a competitive advantage during choice in favor of the encoded action. Such structural encoding also explains two other major defining features of an individual drug preference, its stability over time and its resistance to change.

## INTRODUCTION

Cocaine is the second most used illegal drug in Europe (EMCDDA, 2015). Among people who initiate cocaine use, a substantial number escalate their drug consumption and eventually transition to a state of addiction (i.e., about 10-20%) (Anthony, 2002). During the transition to addiction, cocaine users spend increasingly more time and effort in drug-related activities, often at the expense of other nondrug-related activities that are generally more socially valued. Most of the negative consequences associated with cocaine addiction can in fact be traced to this competitive interference between drug-related versus nondrug-related activities (Martin et al., 2014). This harmful allocation of behavior in favor of drug use has been hypothesized to reflect functional alterations in the basic brain mechanisms underlying decision-making, choice and/or preference in addicted individuals (Volkow and Fowler, 2000, Redish et al., 2008).

This general hypothesis is consistent with previous neuroimaging research in affected individuals showing that cocaine addiction is associated with both structural and functional changes in different prefrontal brain regions and, most notably, in the orbitofrontal cortex (OFC) (Goldstein and Volkow, 2011). This prefrontal region which is also altered in other compulsions (Chamberlain et al., 2008, Milad and Rauch, 2012) has been implicated in many different facets of decision-making and action selection in both human and nonhuman animals, including expected reward value during choice (Tremblay and Schultz, 1999, Padoa-Schioppa and Assad, 2006, FitzGerald et al., 2009), relative preferences for different nondrug rewards or rewarded-actions (Tremblay and Schultz, 1999, Wallis and Miller, 2003, Roesch et al., 2006), and initiation of reward-seeking actions themselves (Homayoun and Moghaddam, 2006, Guillem et al., 2010, Moorman and Aston-Jones, 2014). The information encoded in the OFC probably influences choice via a top-down control of neuronal activity in downstream brain regions involved in action selection and reinforcement learning, such as the striatum and the ventral tegmental area (Gabbott et al., 2005, Schilman et al., 2008). The OFC hypothesis of cocaine addiction is also consistent with evidence for changes in OFC neuronal activity in rats with a history of cocaine self-administration (Lucantonio et al., 2014). However, how exactly OFC neuronal activity encodes choice between cocaine-rewarded and nondrug-rewarded actions, and eventually allocation of behavior in favor of cocaine use in addicted individuals remains unknown.

To begin to address this question, we recorded *in vivo* firing activity of OFC neurons from rats as they chose between a cocaine-rewarded action and an alternative action rewarded by water sweetened with saccharin (Figure 1a). Typically, when given a choice between these different actions, most individual rats preferentially choose the one associated with the nondrug alternative (Lenoir et al., 2007, Cantin et al., 2010, Kerstetter et al., 2012, Tunstall and Kearns, 2014, Tunstall et al., 2014). Only some individual rats (i.e., about 15%) preferentially pursue the action that leads to cocaine at the expense of the other action and despite potentially harmful opportunity costs. This pattern of individual preferences has been observed at different doses of cocaine and even after long histories of cocaine self-administration. Our general hypothesis is that the subgroup of drug-preferring rats would model the subgroup of vulnerable humans who become addicted to cocaine (Ahmed, 2010, Ahmed et al., 2013). Thus, recording firing activity of OFC neurons during choice in drug-preferring *versus* nondrug-preferring rats should provide important insight on how these neurons encode cocaine preference in addicted individuals, and, more generally, on the cortical substrates of individual preferences.

**Figure 1.**
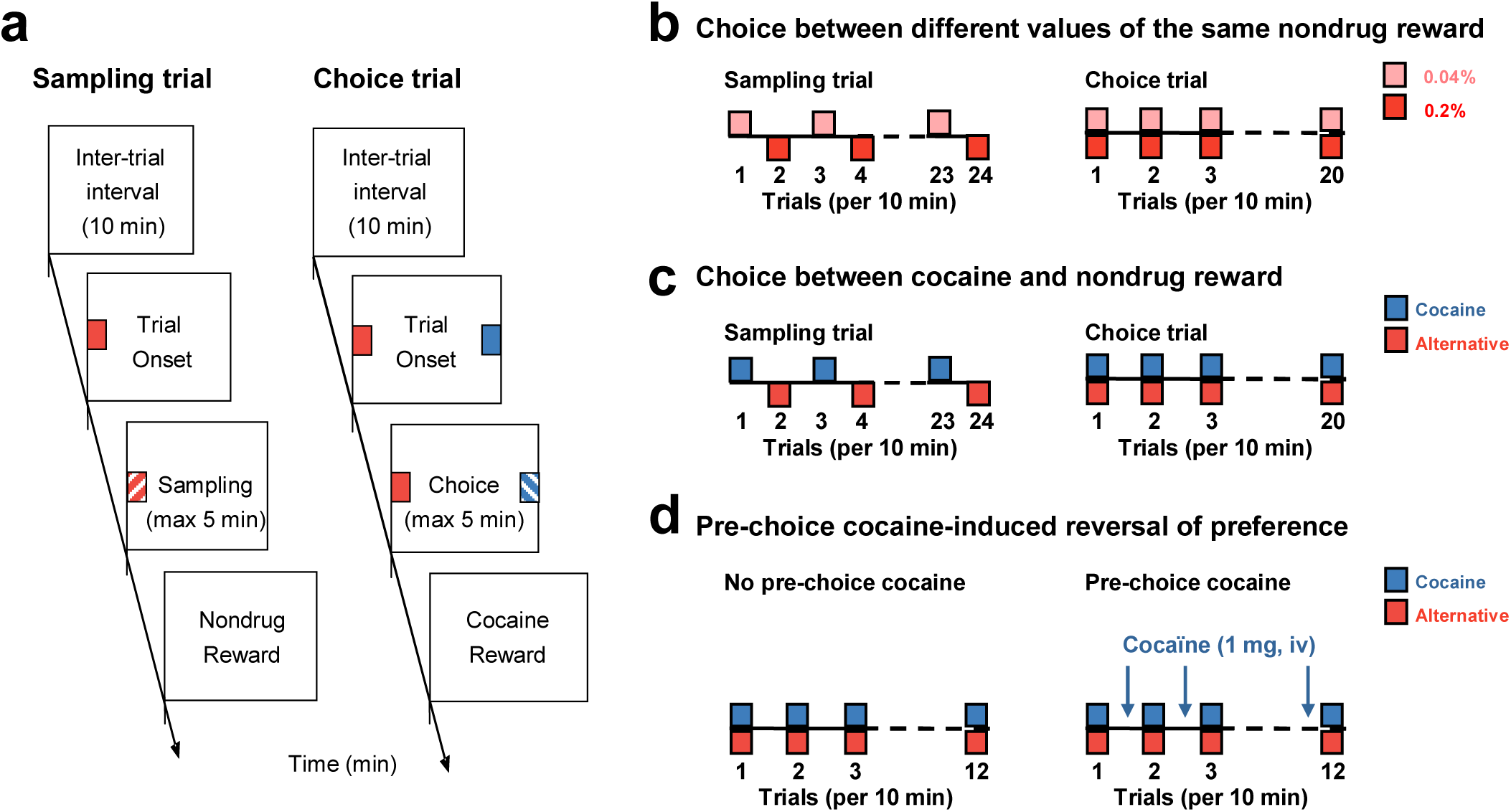
General task design and specific choice procedures. In all cases, the discrete-trials choice procedure was divided into 2 successive phases, first sampling trials followed by choice trials. (**a**) Schematic description of a sampling trial and a choice trial. Rectangles represent different successive task events within each type of trial (i.e., inter-trial interval; trial onset; sampling or choice; reward delivery). (**b,c,d**) Specific discrete-trials choice procedures used in different experiments. (**b**) Choice between two actions rewarded by different values (indicated by different shades of pink) of the same nondrug reward. One action was rewarded by 20-s access to 0.04% saccharin (lighter pink), the other by 20-s access to 0.2% saccharin (darker pink). (**c**) Choice between a cocaine-rewarded (0.25 mg, i.v.) and an alternative, nondrug-rewarded action (20-s access to 0.2% saccharin). (**d**) Choice between a cocaine-rewarded and an alternative, nondrug-rewarded action as in (**c**) but in two different conditions, without (first 12 choice trials) or with pre-choice, non-contingent administration of cocaine (1 mg, i.v.) (last 12 choice trials). Each pre-choice dose of cocaine was administered 5 min before onset of choice trials.

## RESULTS

In the experiments described below, *in vivo* recording of neuronal activity from the OFC was conducted during a single testing session after prior stabilization of preference between two rewarded actions: pressing one of two retractable levers positioned either in the middle of the right (lever R) or the left (lever L) wall of the operant chamber (see Experimental Procedures; Figure 1). Depending on the experimental goal, these two actions were associated with the same or different rewards (Figures 1b-c; see also below). Briefly, the testing session which was similar to previous training sessions consisted of a succession of discrete trials, spaced 10 min apart, and distributed into two main periods, sampling followed by choice (Figure 1a). During sampling or forced choice trials, only one action was available at a time and in alternation with the other action, thereby forcing rats to engage in that action to sample the corresponding reward. During free choice trials, both actions were available simultaneously for choice. Onset of all trials was signaled by the motor noise generated by the automatic insertion of one (sampling period) or two levers (choice period) in the operant chamber and lasted until action completion (i.e., 1 lever press response) or 5 min has elapsed. End of trials was signaled by the automatic retraction of the available lever(s). Previous research has shown that no other cues are needed to signal onset of sampling or choice trials in this procedure (Lenoir et al., 2013).

This unique design allowed us, first, to reliably identify during sampling trials OFC neurons that encode specific aspects of each rewarded action and, then, to follow their differential firing activity during subsequent choice trials (see Experimental Procedures). Though we analyzed neuronal activity during several relevant behavioral events during sampling trials (e.g., conditioned response to automatic lever insertion; reward delivery itself), we primarily based our functional categorization on OFC neurons that encoded the initiation of rewarded actions. This was largely because in all experiments, these neurons were by far the most prevalent in the OFC, a finding consistent with previous research in rats trained to press a lever to obtain a nondrug reward (Kravitz and Peoples, 2008, Moorman and Aston-Jones, 2014). We recorded a total of 439 putative pyramidal neurons in the OFC during sampling in all experiments. These neurons were exclusively located in the lateral (LO; *n* = 279) and ventral (VO; *n* = 160) parts of the OFC (Figure S1). Of these putative pyramidal neurons, 44% encoded one action and/or the other (Supplementary Table 1). In comparison, only 28% of OFC neurons responded to the automatic insertion of the levers during sampling trials. Briefly, a neuron was considered as encoding the initiation of a given action when its firing activity phasically increased and peaked shortly *before* the onset of the action itself (see Results below). The exact role of this phasic firing activity in action initiation is difficult to establish with certainty but we can at least rule out the following other possible roles: motor action itself since it peaked before the performance of the action; degree of confidence that the forthcoming action will be rewarded since each response was rewarded with certainty; conditioned approach to automatic lever insertion since it systematically occurred after lever insertion and/or bouts of locomotion toward the inserted lever (see below). Regardless of the exact information conveyed, however, we called for convenience OFC neurons showing this pre-action phasic activity “action-coding neuron” and used it as a neuromarker to categorize neurons in different groups as a function of the coded action (e.g., cocaine action-versus alternative action-coding neurons).

In a first, control experiment, we checked for any pre-existing bias in behavior and/or in OFC neuronal activity between the actions on lever L or R. To this end, we recorded OFC neuronal activity in a group of rats (*n* = 4) that responded on alternate sampling trials on either lever to obtain the same nondrug reward: 20-s access to water sweetened with 0.2% saccharin (Online Methods). There were no choice trials in this preliminary control experiment. As expected, rats showed no initial difference in responding between lever R or L (Figure 2a). They sampled each lever equally (11.8 ± 0.3 and 11.8 ± 0.3) and with the same latency (6.6 ± 1.9 s and 6.7 ±1.9 s). 45% of neurons recorded in the OFC of these rats phasically encoded action on lever L and/or action on lever R (Figure 2b). Their firing activity phasically increased and peaked shortly before action onset and after termination of locomotion toward the inserted lever (Figure 2b), the latter being not associated with increased firing activity (Figure S2a). Most of these action-coding neurons were nonselective (63%) as they encoded both actions indiscriminately. Only a minority selectively encoded one or the other action but there was no difference in proportion between these two populations of selective neurons (i.e., 16 versus 21%) (Figure 2c). In addition, indexes of action selectivity of all recorded neurons were equally distributed around 0 (*P* = 0.33; Wilcoxon signed-rank test), suggesting that there was no bias in firing activity between these two actions at the population level (Figure 2d). Overall, these findings clearly rule out any preexisting lever side bias in our procedure.

**Figure 2.**
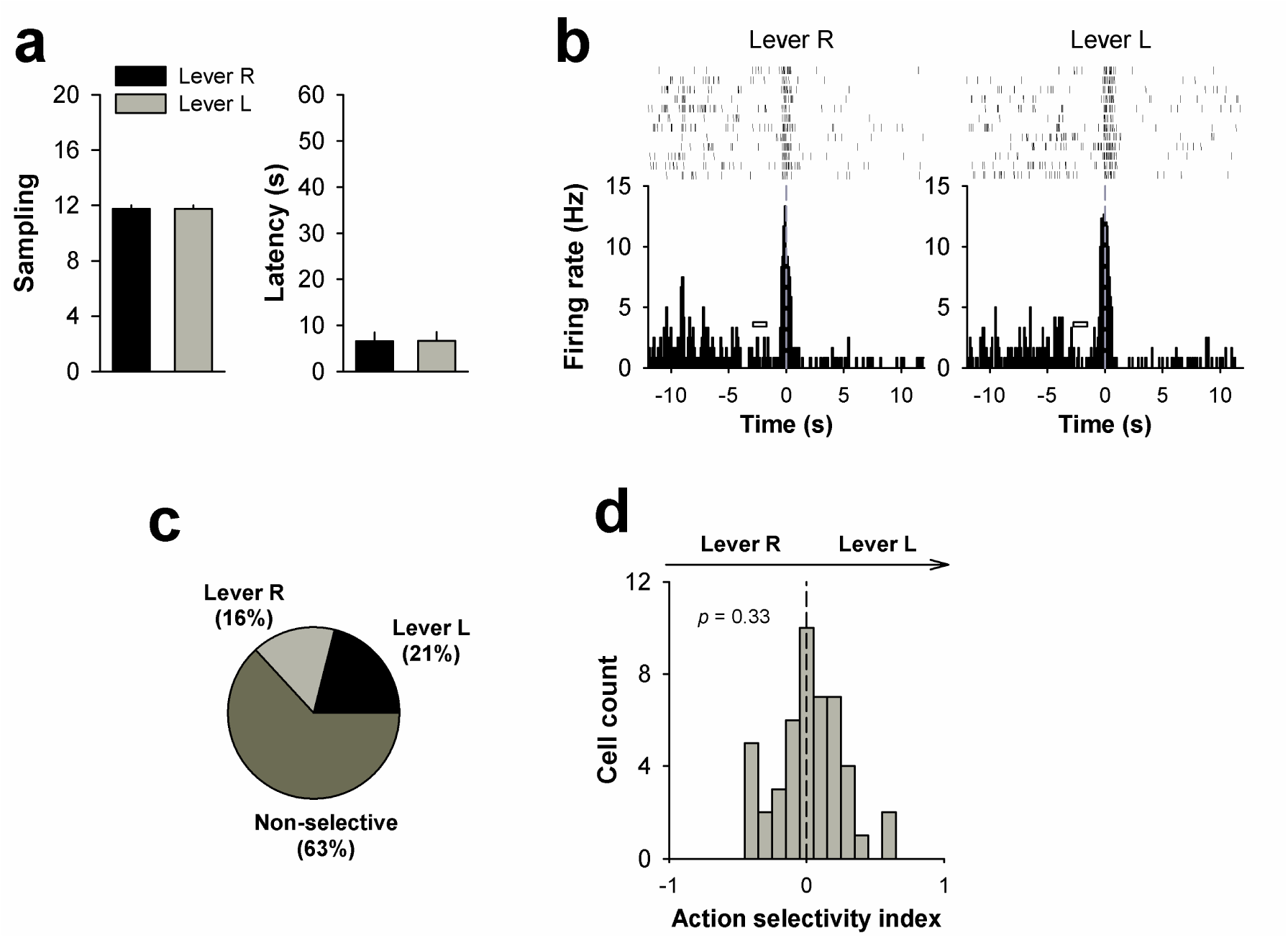
OFC neuronal encoding of two actions equally rewarded by the same nondrug reward. (**a**) Mean number (± s.e.m.) of completed sampling trials and mean sampling latency (± s.e.m.) as a function of lever side (R, right; L, left), (**b**) Examples of selective L or R action-coding neurons. Each perievent histogram shows the firing rate of a single neuron plotted as a function of time, with 0 corresponding to the onset of each lever press action. The small horizontal rectangle on the left of each firing peak indicates the period when bouts of locomotion toward the corresponding lever were recorded. (**c**) Pie chart representation of the distribution of OFC action-coding neurons into 3 categories: selective to action on lever R, selective to action on lever L, and non-selective. (**d**) Distribution of neuronal indexes of action selectivity. This distribution was centered on 0, indicating that the population of OFC neurons as a whole fired equally during both actions (see Methods, for additional information).

We next sought to measure how OFC neurons encode two actions rewarded by different values of the same nondrug reward (i.e., water sweetened with different concentrations of saccharin) and how these neurons fire during choice and preference between these two actions (Figure 1b). One action was associated with 20-s access to 0.2% saccharin and the other action with 20-s access to a 5-fold lower concentration of saccharin, 0.04%. As expected, during sampling, all rats engaged more (*F*_(1,6)_ = 14.1; *P* < 0.05) and faster (*F*_(1,6)_ = 16.9; *P <* 0.01) in the action rewarded by the highest concentration of saccharin (Figure 3a), suggesting that it had acquired a greater value. This was directly confirmed during choice as all rats expressed a marked preference for 0.2 over 0.04% of saccharin (*F*_(1,6)_ = 239; *P* < 0.001) (Figures 3a,b).

**Figure 3.**
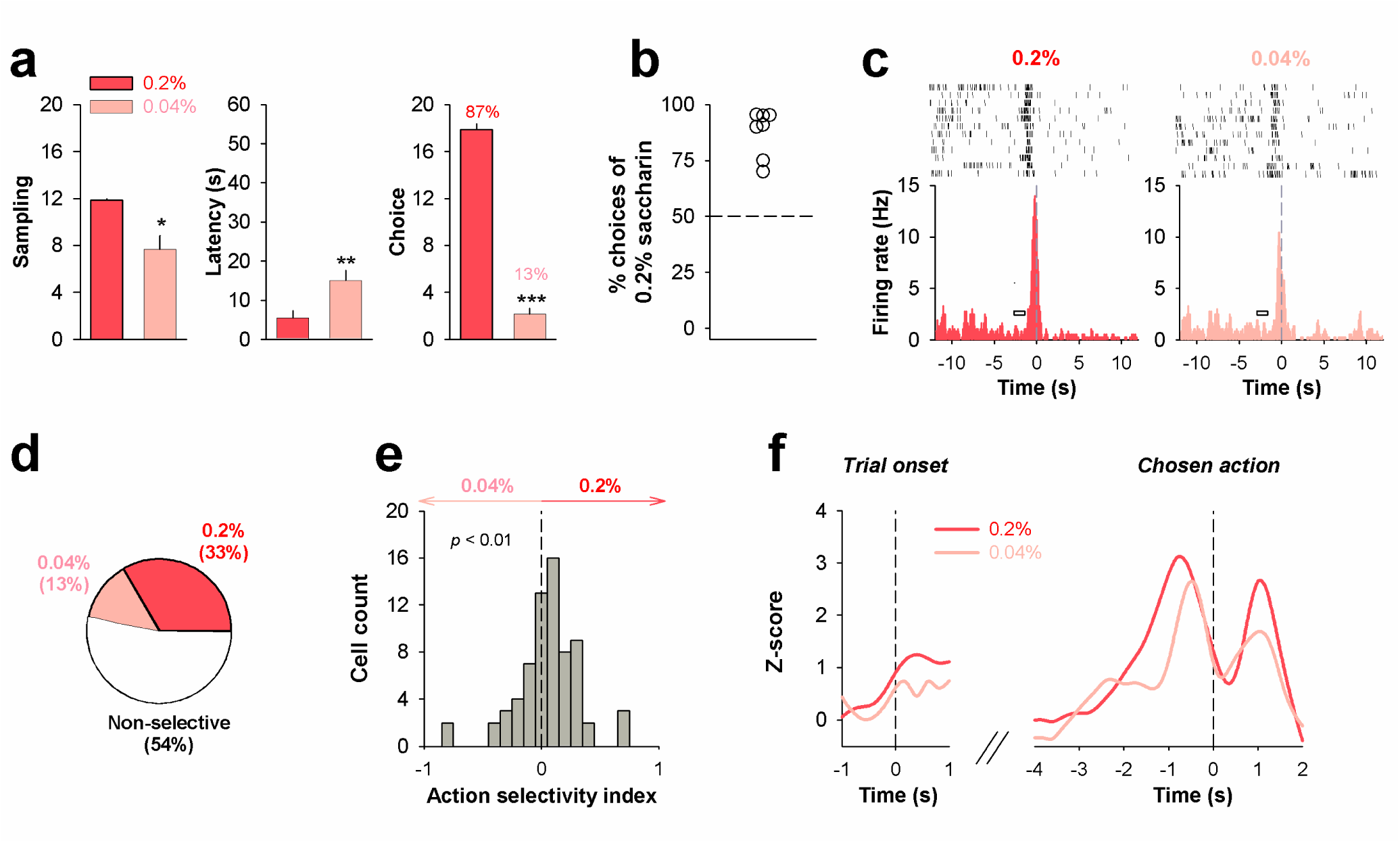
OFC neuronal encoding of preference between two actions rewarded by different values of the same nondrug reward. (**a**) Mean number of sampling (± s.e.m.), mean sampling latency (± s.e.m.), and mean number of choice (± s.e.m.) as a function the concentration of saccharin associated with each action. \**p* < 0.05, \*\**p* < 0.01 and \*\*\**p* < 0.001, different from 0.2%. (**b**) Scatter plot of individual % choices of the action rewarded by 0.2% saccharin. (**c**) Examples of selective action-coding neurons. Each perievent histogram shows the firing rate of a single neuron plotted as a function of time, with 0 corresponding to the onset of each lever press action. The small horizontal rectangle on the left of each firing peak indicates the period when bouts of locomotion toward the corresponding lever were recorded. (**d**) Pie chart representation of the distribution of OFC action-coding neurons into 3 categories: selective to the action rewarded by 0.2% saccharin, selective to the action rewarded by 0.04% saccharin, and non-selective. (**e**) Distribution of neuronal indexes of action selectivity. The distribution was significantly shifted to the right, indicating that the population of OFC neurons as a whole fired more during the preferred action. (**f**) Mean normalized firing rate (z-scores smoothed with a Gaussian filter) of the two groups of selective action-coding neurons at onset of choice trials and during actual choice of the preferred action.

At the neuronal level, we identified 3 main categories of action-coding neurons in the OFC during sampling trials: neurons that selectively encoded the preferred action (i.e., rewarded by 0.2% saccharin), neurons that selectively encoded the non-preferred action (i.e., rewarded by 0.04% saccharin) (Figure 3c), and, finally, non-selective neurons that encoded indistinctly both actions (not shown). In all cases, firing activity phasically increased and peaked shortly before the onset of the coded action and after termination of locomotion toward the inserted lever (Figure 3c), the latter being not associated once again with increased firing activity (Figure S2b). About half of action-coding neurons (54%) were nonselective while the other half (46%) were selective for one action or the other (Figure 3d). Importantly, among selective action-coding neurons, the proportion encoding the preferred action (33%) was about 3 times larger than that encoding the non-preferred one (13%) (Z = 2.7; *P* < 0.01) (Figure 3d). Moreover, at the population level, the distribution of action selectivity indexes was shifted significantly to the right (*P* < 0.01; Wilcoxon signed-rank test), indicating that OFC neuronal activity is dominated by neurons encoding the preferred action (Figure 3e). Thus, there was a good correspondence between the relative proportion of OFC neurons selectively encoding a nondrug-rewarded action among other options and the preference for that action.

Once OFC selective action-coding neurons were identified during sampling trials, we followed and analyzed their firing activity during subsequent choice trials where rats chose the preferred action. By necessity, choice trials where rats chose the non-preferred action were too rare to be analyzed. Specifically, the activity of selective action-coding neurons was aligned to two major pre-choice events: the onset of choice trials signaled by the automatic insertion of the levers and the onset of the preferred action itself. Rapidly after onset of choice trials and well before the preferred action (i.e., within less than 1 sec), neurons encoding the preferred action slightly increased their firing activity above that of neurons encoding the non-preferred action (Figure 3f). Surprisingly, firing activity of both groups of neurons also increased and peaked shortly before the preferred action. However, the increase was faster and higher in preferred action-coding neurons than in non-preferred action-coding neurons. Overall, these findings strongly indicate that preference for a nondrug-rewarded action is somehow determined by the relative size and pre-choice activity of the neuronal population encoding that action in the OFC – a conclusion generally consistent with previous similar research (Tremblay and Schultz, 1999, Wallis and Miller, 2003).

Based on this solid experimental foundation, we next assessed how OFC neurons encode choice and preference between two actions rewarded by two distinct kinds of reward, a drug and a nondrug reward. Animals (*n* = 10) were trained to sample and choose between one action rewarded by an intravenous infusion of cocaine (0.25 mg) and the other rewarded by water sweetened with 0.2% saccharin (Figure 1c). Consistent with previous research (Lenoir et al., 2007, Cantin et al., 2010), rats engaged significantly more (11.6 ± 0.3 and 8.6 ± 0.8; *F*_(1,9)_ = 12.3, *P* < 0.01) and faster (11.3 ± 2.6 and 43.1 ± 5.1 s; *F*_(1,9)_ = 25.4, *P* < 0.001) in the nondrug-rewarded action than in the cocaine-rewarded action, indicating that the former had acquired more value than the latter (Figure 4a). This was confirmed during choice by an overall preference for the nondrug action over the cocaine action (*F*_(1,9)_ = 5.3, *P* < 0.05) (Figure 4a). As expected, however, though many rats preferred the nondrug action (*n* = 5, red circles, % choices of the nondrug action > 65%), few others preferred the cocaine action (*n* = 2, blue circles, % cocaine choices > 65%) or were indifferent (*n* = 3, white circles) (Figures 4b,c).

**Figure 4.**
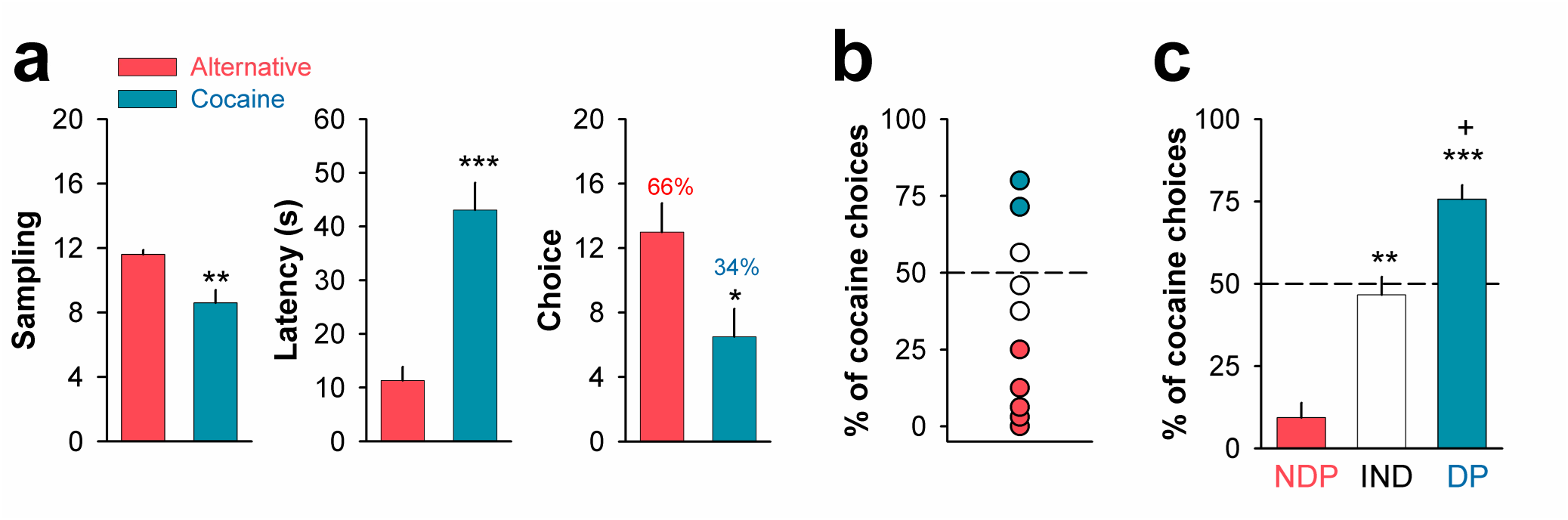
Variation in individual drug preferences. (**a**)Mean number of sampling (± s.e.m.), mean sampling latency (± s.e.m.), and mean number of choice (± s.e.m.) as a function of the type of reward associated with each action. \**p* < 0.05 and \*\**p* < 0.01, different from the nondrug alternative. (**b**) Scatter plot of individual % cocaine choices reveals 3 subgroups of rats: nondrug-preferring or NDP rats (red circles, *n* = 5), drug-preferring or DP rats (blue circles, *n* = 2), and indifferent or IND rats (white circles, *n* = 3). (c) Mean % cocaine choices (± s.e.m.) as a function of the subgroup of rats. \*\**p* < 0.01 and \*\*\**p* < 0.001, different from NDP rats. ^+^*p* < 0.05, different from IND rats.

We therefore analyzed OFC neuronal activity as a function of these different subgroups. First and irrespective of individual choice behavior, we again identified 3 main categories of action-coding neurons in the OFC during sampling trials: neurons that selectively encoded the cocaine action, neurons that selectively encoded the nondrug alternative (Figure 5a) and, finally, neurons that non-selectively encoded both actions indiscriminately (not shown). In all cases, firing activity phasically increased and peaked shortly before the onset of the coded action and after termination of locomotion toward the inserted lever (Figure 5a), the latter being not associated with increased firing activity (Figure S2c). Second and most importantly, we found that the relative proportion of cocaine action-versus alternative action-coding neurons in the OFC was positively correlated with individual drug preferences during choice trials (Figure 5b). In nondrug-preferring (NDP) rats, the proportion of alternative action-coding neurons was much higher than that of cocaine action-coding neurons (53 versus 22%; Z = 2.6; *P* < 0.01) (Figure 5c). In indifferent (IND) rats, alternative action-coding neurons tended also to be more prevalent than cocaine action-coding neurons but this difference was no longer significant (44 versus 33%, Z = 0.7; NS) (Figure 5d). In contrast, in drug-preferring (DP) rats, the opposite trend was true: the proportion of cocaine action-coding neurons was slightly, albeit non-significantly, higher than that of alternative action-coding neurons (38 versus 31%; Z = 0.6; NS) (Figure 5e). Importantly, this inversion in the OFC neuronal representation of both actions in DP rats was not due to an increased recruitment of cocaine action-coding neurons from the pool of non-selective neurons compared to NDP rats. Instead it resulted almost equally from an increase in cocaine action-coding neurons (from 22% to 38%; Z = 1.6; *P* < 0.05) and a decrease in alternative action-coding neurons (from 53 versus 31%; Z = 1.7; *P* < 0.05), thereby suggesting a functional competition between these two groups of OFC neurons. At the population level, this neuronal inversion was reflected by an opposite shift in the distribution of action selectivity indexes, either to the right in NDP rats (*P* < 0.05; Wilcoxon signed-rank test) (Figure 5f) or to the left in DP rats (*P* < 0.05; Wilcoxon signed-rank test) (Figure 5h). In IND rats, the distribution of action selectivity indexes centered on 0 (*P* = 0.25; Wilcoxon signed-rank test) (Figure 5g). Thus, as with preference between two nondrug-rewarded actions (see above), there was a good fit between the relative proportion of OFC neurons selectively encoding a drug-versus nondrug-rewarded action and preference for that action.

**Figure 5.**
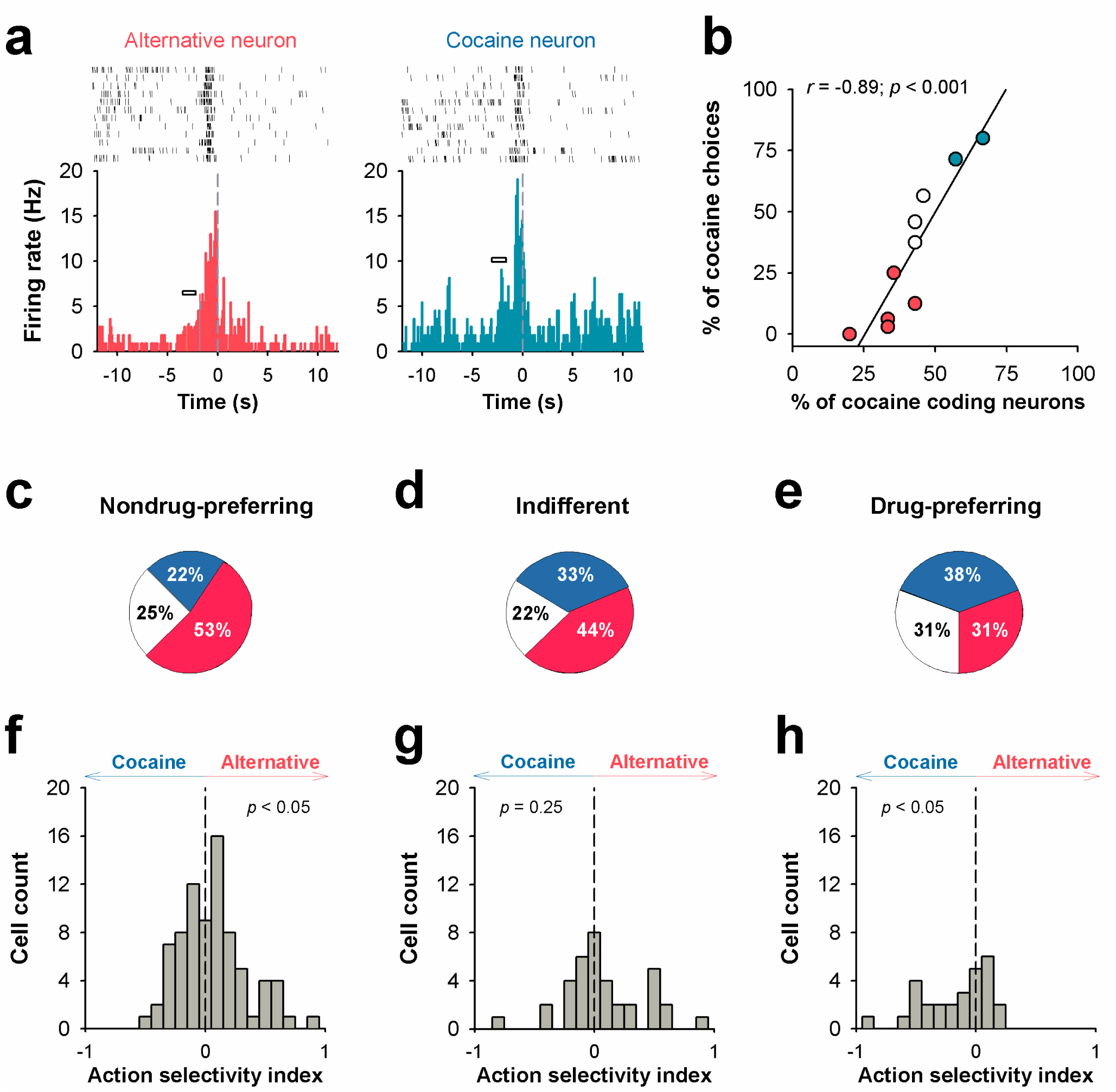
OFC neuronal encoding of individual preferences between a cocaine-rewarded and an alternative nondrug-rewarded action, (**a**) Examples of selective action-coding neurons. Each perievent histogram shows the firing rate of a single neuron plotted as a function of time, with 0 corresponding to the onset of each lever press action. The small horizontal rectangle on the left of each firing peak indicates the period when bouts of locomotion toward the corresponding lever were recorded, (**b**) Pearson correlation between individual % cocaine choices and individual % cocaine action-coding neurons, (**c-e**) Pie chart representation of the distribution of OFC neurons into selective and non-selective action-coding categories in NDP rats (**c**), IND rats (**d**) and DP rats (**e**). (**f-h**) Distribution of neuronal indexes of action selectivity in NDP rats (**f**), IND rats (**g**) and DP rats (**h**) (see Results, for additional information).

Once selective action-coding neurons were identified in the OFC during sampling trials, we followed and analyzed their pre-choice firing activity during subsequent choice trials (i.e., at both trial and action onset) as a function of individual preferences (Figures 6a-c). As explained above, for each subgroup, except for indifferent rats, analysis was restricted to choice trials where rats chose their preferred action (i.e., the cocaine action in DP rats and the alternative action in NDP rats). Regardless of the preferred action (be it that rewarded by cocaine or the nondrug alternative), neurons encoding the preferred action fired more than neurons encoding the non-preferred action both shortly after the onset of choice trials (i.e., within 1 sec) and shortly before the preferred action itself (NDP rats: Figure 6a; DP rats: Figure 6c). Thus, preference for a cocaine-or a nondrug alternative-rewarded action was somehow determined by the relative size and pre-choice activity of the neuronal population encoding that action in the OFC.

Interestingly, in IND rats which chose about equally frequently the cocaine action or its alternative during choice trials (11.0 ± 1.2 versus 12.7 ± 1.5, respectively), neurons encoding the cocaine action tended to fire more than neurons encoding the alternative action on trials where they chose the cocaine action (Figure 6b1) and, vice versa, on trials where they chose the alternative action (Figure 6b2). This outcome shows that the state of differential pre-choice activity between OFC neurons that encode two equally-preferred actions reflects which action will be chosen on a given trial. It also reveals that indifference between two actions, as opposed to preference for one action, results from an inability of one of the two groups of action-coding neurons to systematically fire more than the other before choice, presumably because of the approximately equal size of both groups of neurons (Figure 5d).

**Figure 6.**
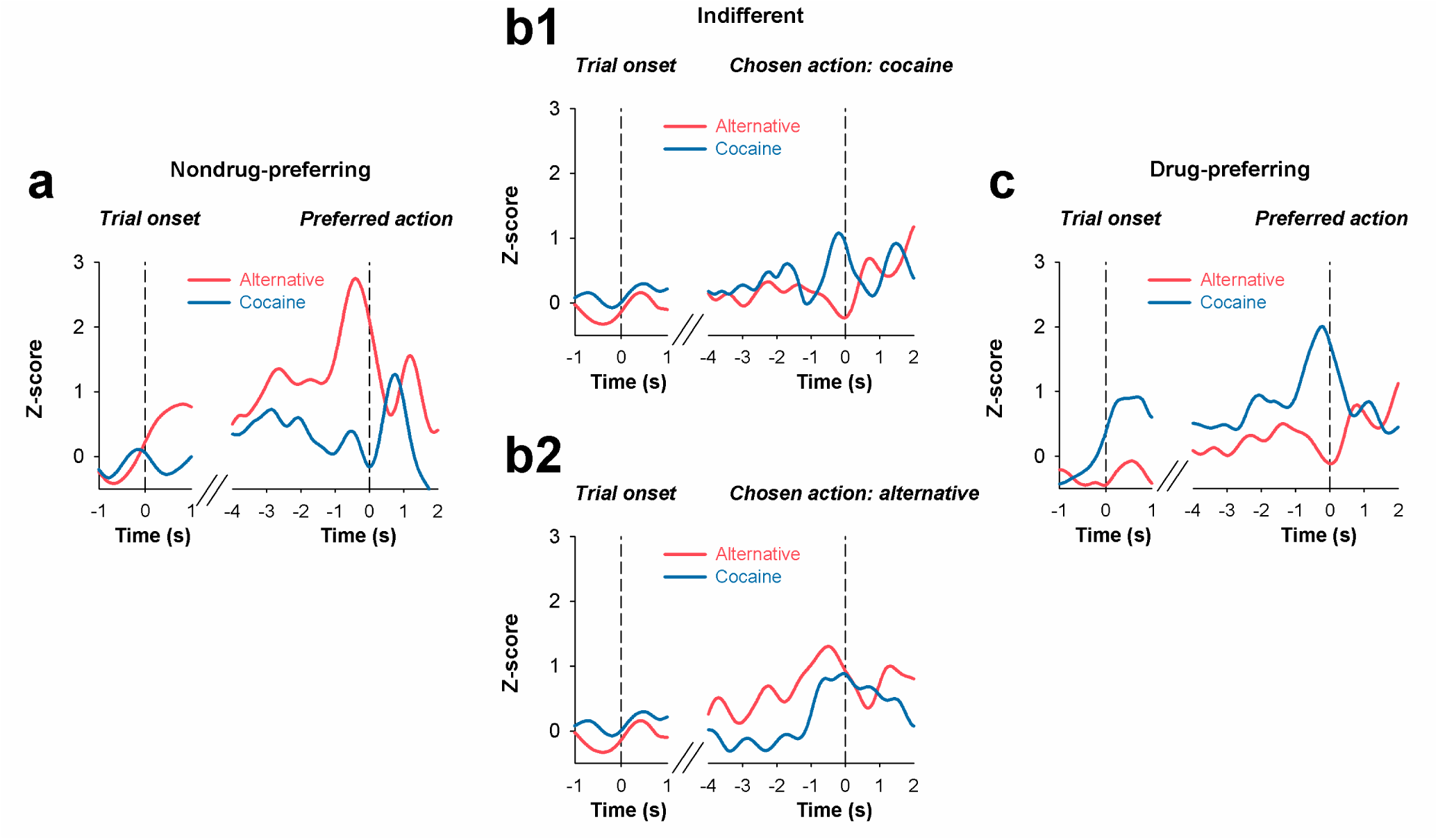
OFC firing activity before and during choice of preferred action as a function of individual drug preferences, (**a-c**) Mean normalized firing rate (z-scores smoothed with a Gaussian filter) of the two groups of selective action-coding neurons at onset of choice trials and during actual choice of the preferred or equally preferred action in NDP rats (**a**), IND rats (**b1,2**) and DP rats (**c**) (see Results, for additional information).

Finally, to further probe the role of differences in pre-choice activity between cocaine action-and alternative action-coding neurons, we recorded OFC neuronal activity in a separate group of NDP rats (*n* = 5) during reversal of preference to cocaine. Preference reversal was induced by passively administering cocaine before each choice trial (Figure 1d). Previous research has shown that this pharmacological intervention reliably and immediately shifts choice to cocaine almost exclusively and in nearly all NDP rats (Vandaele et al., 2016). Importantly, this shift is not due to an increased motivation for drug use but instead to a suppressed motivation to engage in the preferred action rewarded by sweet water induced by the anorexic effects of cocaine (Balopole et al., 1979, Cooper and van der Hoek, 1993). Therefore, we hypothesized that passive, pre-choice administration of cocaine shifts preference to cocaine by decreasing pre-choice firing activity of alternative action-coding neurons, with little or no effect on cocaine action-coding neurons.

As expected, compared to the control condition where rats do not receive pre-choice cocaine, passive, pre-choice administration of cocaine shifted preference from the alternative action to the cocaine action almost exclusively in all rats (*F*_(1,4)_ = 23.8; *P* < 0.01) (Figure 7a).

**Figure 7.**
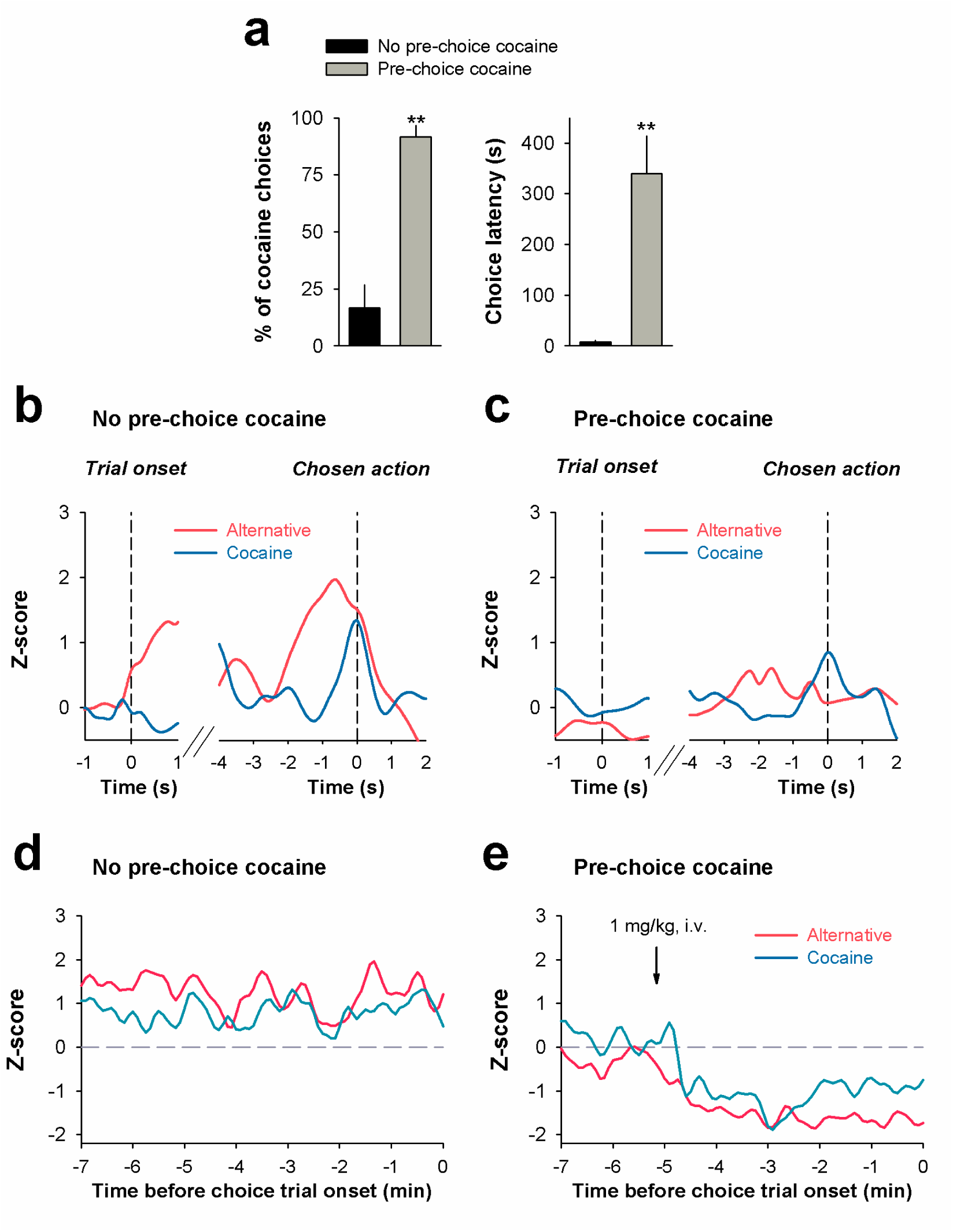
OFC neuronal encoding of preference reversal in NDP rats. (**a**) Pre-choice cocaine-induced shift of preference to cocaine. Mean cocaine choice (± s.e.m.) and mean choice latency (± s.e.m.) without (first 12 trials) or with pre-choice administration of cocaine (last 12 trials). Each pre-choice dose of cocaine was administered 5 min before onset of choice trials. \*\**p* < 0.01, different from the “no pre-choice cocaine” condition. (**b,c**) Mean normalized firing rate (z-scores smoothed with a Gaussian filter) of the two groups of selective action-coding neurons at onset of choice trials and during actual choice of the preferred action in two conditions: without (**b**) or with pre-choice administration of cocaine (**c**). (**d,e**) Cocaine induces suppression of OFC firing activity. Mean normalized firing rate (z-scores smoothed with a Gaussian filter) during inter-trial intervals during choice trials without (**d**) or with pre-choice administration of cocaine (**e**). The downward arrow indicates when pre-choice cocaine was administered (i.e., 5 before the onset of choice trials).

This shift of preference was associated with a dramatic increase in choice latency (*F*_(1,4)_ = 18.4; *P* < 0.01), presumably due to recovery from drug satiation. Thus, though rats could choose to engage in their normally preferred action while drug-sated, they nevertheless did not, confirming that their motivation for this action was suppressed. As hypothesized, the shift of preference induced by pre-choice cocaine was associated with an inversion in pre-choice neuronal activity between alternative action-and cocaine action-coding neurons. After pre-choice cocaine, the former fired less than the latter – a difference opposite to that normally seen in NDP rats during baseline (Figures 7b versus 7c). Interestingly, this effect seems to be mainly due to a suppression of activity of alternative action-coding neurons rather than to a facilitation of activity of cocaine action-coding neurons. This conclusion was confirmed by an analysis of the effects of pre-choice cocaine on OFC neuronal activity during the inter-trial intervals before choice trials. During the control condition (i.e., no pre-choice cocaine), firing activity of alternative action-coding neurons tended to be systematically higher than that of cocaine action-coding neurons (Figure 7d). In contrast, after passive, pre-choice administration of cocaine, this difference in inter-trial neuronal activity was inversed in favor of cocaine action-coding neurons (Figure 7e). Though pre-choice cocaine suppressed both groups of action-coding neurons, it suppressed more intensely alternative action-coding neurons than cocaine action-coding neurons (Figures 7b,c). As a result of this differential suppression, the latter remained more active than the former until onset of choice trials, thereby probably inducing the reversal of preference to cocaine observed at the behavioral level. Taken together, these findings strongly suggest that differential pre-choice activity of cocaine action-versus alternative action-coding neurons plays a causal role in which action will be chosen during choice.

## DISCUSSION

Here we identified in the OFC a mechanism that explains individual preferences during choice between a cocaine-rewarded action and an alternative, nondrug-rewarded action. Most notably, we found that the relative size and differential pre-choice activity of the populations of OFC neurons that selectively encode initiation of these actions seem to determine individual drug preferences. Previous research in other brain regions, downstream to the OFC, has also shown that different, non-overlapping populations of neurons can encode cocaine use versus other nondrug actions, such as juice consumption (Bowman et al., 1996, Opris et al., 2009, Cameron and Carelli, 2012). However, it is the first time that a relationship is observed between an individual preference for a specific action, notably when it is oriented toward cocaine, and the relative size and differential pre-choice activity of the population of neurons that selectively encodes the initiation of that action.

Overall, this study suggests a heuristic framework to conceptualize how OFC neurons encode individual preferences in both non-addicted and cocaine-addicted rats. An individual preference for a specific rewarded action appears to be determined by the larger size of the neuronal population that encodes that action in the OFC, relative to other, smaller action-coding populations. A large size is a relatively stable, structural feature that may confer to a population or ensemble of OFC neurons a systematic advantage if it is in competition with other neighbor populations during choice-making (e.g., to gain access to and control a same downstream mechanism of action selection) (Newsome et al., 1989, Samejima et al., 2005, Kim et al., 2009, Ito and Doya, 2015). Though the underlying mechanisms of this structural advantage need to be defined, it may explain why in both drug-and nondrug-preferring rats, preferred action-coding neurons fire more than non-preferred action-coding neurons during almost all choice trials, thereby biasing choice toward the preferred action. Conversely, this may also explain why in indifferent rats where the two populations of action-coding neurons were roughly of equal size, both populations fired more than the other about equally frequently on alternate choice trials.

The encoding of individual preferences in a structural brain difference also provides a simple explanation for why such preferences are stable over time and difficult to change, particularly when oriented toward cocaine use. A key question for future research will be to understand how these different populations of action-coding neurons emerge during preference acquisition and fixation, and how they acquire their different sizes. As shown in the present study, it seems that the pool of OFC neurons allocated to selective action coding is rather limited and cannot be extended by recruiting additional neurons from the pool of OFC neurons dedicated to non-selective action coding. As a result, the distribution of neurons into different populations according to the encoded action is probably also competitive. Understanding the mechanisms underlying this competitive neuronal distribution will represent a breakthrough in the field, especially if it helps to explain the development of preference for cocaine use in addicted individuals and if it can generalize to other drug addictions. This future research will eventually be facilitated by the development of adapted in vivo deep brain calcium imaging methods to record activity of hundreds of OFC neurons in the same individual animals (Pinto and Dan, 2015, Ziv and Ghosh, 2015), coupled with theoretical models of neuronal population competition (Gold and Shadlen, 2002, Rangel et al., 2008). Finally, this study suggests that stable reversal of individual drug preferences will eventually require developing selective interventions that change the relative size of drug action-coding neuronal populations in the OFC.

## EXPERIMENTAL PROCEDURES

### Subjects

A total of 26 male Wistar rats (275–300 g) (Charles River Laboratories, L’Arbresle, France) were individually housed in cages containing enrichment materials (a nylon bone and a cardboard tunnel; Plexx BV, The Netherlands) under a 12 h light/dark cycle, with ad libitum access to food and water. All experiments were carried out in accordance with standard ethical guidelines (European Communities Council Directive 86/609/EEC) and approved by the committee on Animal Health and Care of Institut National de la Santé et de la Recherche (Agreement A5012052).

### Surgeries

Animals were anesthetized with a mixture of ketamine (100 mg/kg, i.p., Bayer Pharma, Lyon, France) and xylazine (15 mg/kg, i.p., Merial, Lyon, France) and surgically prepared with an indwelling silastic catheter in the right jugular vein (Dow Corning Corporation, Michigan, USA). Then, under isofluorane anesthesia, arrays of 16 teflon-coated stainless steel microwires (MicroProbes Inc, Gaithersburg, MD) were implanted unilaterally in the OFC [AP: + 2.5 to + 3.7 mm, ML: 1.5 to 4.5 mm, and DV: - 5.0 mm relative to skull level] as previously described (Guillem et al., 2010). A stainless steel ground wire was also implanted 4 mm into the ipsilateral side of the brain, 5 mm caudal to bregma. After surgery, catheters were flushed daily with 0.2 ml of a sterile antibiotic solution containing heparinized saline (280 IU / ml) and ampicilline (Panpharma, Fougères, France). When a leakage in the catheter was suspected, its patency was checked by an intravenous administration of etomidate (0.75-1 mg/kg, Braun Medical, Boulogne-Billancourt, France), a short-acting non-barbiturate anesthetic. Behavioral testing began 10-14 days after surgery.

### Behavioral procedures

#### Behavioral apparatus

The operant chamber (30 × 40 × 36 cm) used for all behavioral training and testing (Imetronic, Pessac, France) has been described in detail elsewhere (Madsen and Ahmed, 2015). Briefly, the chamber was equipped with 2 automatically retractable levers located on the middle of the left and right walls of the chamber (see Figure 1). These positions increased the discriminability between the levers which were otherwise physically similar while minimizing any spatial preference or bias (see Results). Each lever was equipped with a photobeam device to precisely detect and record lever presses. A white cue light was positioned above each lever. The chamber was also equipped with two drinking cups, each located 6.5 cm on the left of each lever on the same wall and 6 cm above the grid floor. Each drinking cup was connected to a lickometer circuit to record cup licks. Finally, the chamber was also equipped with two computer-controlled syringe pumps connected to different line tubings for drug delivery and/or fluid delivery into the cups, a single-channel liquid swivel (Lomir Biomedical Inc., Quebec, Canada), an electrical commutator (Crist Instrument Inc., Hagerstown, MD), and two pairs of infrared beams to measure horizontal cage crossings.

#### Discrete-trials choice procedure

Before neuronal recording during choice, all rats were first trained on alternate days to respond on lever L or R under a fixed-ratio 1 (FR1 time-out 20s) schedule to obtain different rewards (see below). Then, rats were trained under a discrete-trials choice procedure during several days until stabilization of preference, as described previously (Lenoir et al., 2013) (Figure 1a). During testing, this procedure was modified to include a sufficient number of sampling and choice trials to increase the power to detect statistical changes in neuronal activity. Briefly, each choice session consisted of 44 trials spaced 10 min apart and distributed into two successive phases, first sampling (24 trials), then choice (20 trials). During sampling, the two levers were presented separately on alternate trials, thereby forcing rats to engage in only one rewarded action at a time (e.g., pressing lever R). Thus, during sampling, trial onset was systematically signaled by presentation of only one lever. If rats responded within 5 min on the presented lever, they received the reward associated with that action (see below). Reward delivery was signaled by immediate retraction of the lever and illumination of a 40-s illumination of the cue-light above it. If rats failed to respond within 5 min, the lever retracted and no cue-light or reward was delivered. During choice, the two levers were presented simultaneously on the same trials, thereby allowing rats to choose between the two corresponding rewarded actions (i.e., pressing either lever L or R). Thus, during choice, trial onset was signaled by the simultaneous presentation of both levers. If rats responded within 5 min on either lever, they received the reward associated with that action. Reward delivery was signaled by immediate retraction of both levers and illumination of a 40-s illumination of the cue-light above the chosen lever. If rats failed to respond on either lever within 5 min, both levers retracted and no cue-light or reward was delivered.

### Specific experiments

#### Neuronal activity during performance of two actions rewarded by the same nondrug reward

We recorded OFC neuronal activity in a group of rats (*n* = 4) that responded on alternate sampling trials on lever R or L to obtain the same nondrug reward, that is: 20-s access to water sweetened with 0.2% saccharin delivered in a nearby drinking cup (see Apparatus). This experiment allowed us to detect any initial spatial preference bias in behavior and/or OFC neuronal activity. In theory, if there was a spatial preference bias, then a difference in performance and/or OFC neuronal activity should be observed between the action of pressing lever L and the action of pressing lever R, despite being identically rewarded. Note that rats were not tested for choice trials in this preliminary control experiment.

#### Neuronal activity during choice between two actions rewarded by different values of the same nondrug reward

We recorded neuronal activity in a separate group of rats (*n* = 7) tested under the discrete-trials choice procedure described above. In this experiment, the two actions available (i.e., pressing lever L or R) were rewarded by different values of the same nondrug reward. Specifically, one action was associated with 20-s access to water sweetened with 0.2% saccharin, the other with 20-s access to water sweetened with 0.04% saccharin (Figure 1b). In both cases, sweet water was delivered in a cup nearby the location of the corresponding action (i.e., left or right cup).

#### Neuronal activity during choice between a cocaine-rewarded and a nondrug-rewarded action

We recorded OFC neuronal activity in a separate group of rats (*n* = 10) tested under the discrete-trials choice procedure described above. In this experiment, one action was rewarded by 0.25-mg cocaine delivered intravenously over 5 s and the other by 20-s access to water sweetened with 0.2% saccharin delivered in a nearby cup (Figure 1c). Based on their individual cocaine choices, rats were allocated into one of 3 groups: 1) nondrug-preferring (NDP) rats which represented the majority (cocaine choices < 35%, *n* = 5), 2) drug-preferring (DP) rats (cocaine choices > 65%, *n* = 2), and 3) indifferent (IND) rats (cocaine choices between 35 and 65%, *n* = 3).

#### Neuronal activity during drug-induced reversal of preference

We recently reported that non-contingent cocaine administration (i.e., 1 mg) before each choice trial induced a robust and immediate reversal of preference to cocaine in nondrug-preferring rats (Vandaele et al., 2016). This acute reversal mainly resulted from the direct suppressive effects of cocaine on responding for and intake of sweet water. Thus, when rats were under this negative influence at the moment of choice, they lose interest in the nondrug-rewarded action in favor of the cocaine-rewarded action. Here we studied the neuronal correlates of this reversal of preference to better understand the role of OFC neuronal activity during choice. We recorded OFC neurons in a separate group of rats (*n* = 5) under the same discrete-trials choice procedure described above, except that the last 12 choice trials were each preceded by a pre-choice administration of i.v. cocaine (Figure 1d). No passive administration was programmed before sampling trials. Pre-choice cocaine (1 mg) was not signaled and was delivered non-contingently 5 min before each choice trial. Importantly, the dose of cocaine (0.25 mg) available for self-administration during choice trials remained unchanged.

### Data analysis

#### Neuronal recordings and single unit isolation

Voltage signals from each microwire were recorded, amplified up to 40000X, processed, and digitally captured using commercial hardware and software (OmniPlex, Plexon, Inc, Dallas, TX). Spiking activity was digitized at 40 kHz, bandpass filtered from 250 Hz to 8kHz and captured using commercial hardware and software (Plexon). Behavioral events in the operant task were streamed to the Plexon via TTL pulses delivered from the Imetronic system (Imetronic, Pessac, France) to allow the neuronal data to be accurately synchronized and aligned to these behavioral events. Single units spike sorting was performed off-line: principal component scores were calculated and plotted in a three-dimensional principal component space and clusters containing similar valid waveforms were defined (Offline Sorter, Plexon). The quality of individual-neuron recordings was ensured with the following criteria: <3% of all interspike intervals exhibited by the unit were < 2000 μs, and the average amplitude of the unit waveform was at least three times larger than that of the noise band (>3:1 signal-to-noise). The total data set consisted of 477 OFC single units. These units were classified into putative interneurons and pyramidal neurons according to the waveform spike width and average firing rate, as previously described (Guillem et al., 2010). In the population of recorded single units, only 8% were classified as narrow spiking, putative interneurons (width < 300 μs). In this report, we focus exclusively on the remaining 439 units, a population that included putative pyramidal neurons. Electrophysiological data were analyzed using NeuroExplorer (Plexon) and Matlab (Mathworks, Natick, MA).

#### Neuronal data analysis

Peri-event firing rate histograms (PETHs) were used to analyze neuronal responses to relevant behavioral events (i.e., onset of action during sampling and choice trials, and onset of choice trials which were signaled by the automatic insertion of the levers). Neurons were tested for phasic changes in firing time-locked to specific events by comparing firing rate during specific time windows using a Wilcoxon test. For phasic changes associated with onset of choice trials, firing rates during 1 s before and 1 s after trial onset were compared. For phasic changes associated with action initiation during sampling or choice trials, firing rate during 1 s before and 1 s after action onset (recorded as a beam break) was compared to average firing rate during a 3-s window period recorded 9 s before the action. In each experiment, the percent of responding neurons was determined for each action and compared using Z-test statistics. An index of action selectivity (Ogawa et al., 2013) was calculated for each recorded neuron as follows: (firing rate during action X - firing rate during action Y) / (firing rate during action X + firing rate during action Y). An index equal to 0 indicates that the corresponding neuron has no action selectivity. In contrast, an index greater or lower than 0 indicates that the corresponding neuron fires more or less during action X than during action Y, respectively. The position of the distribution of action selectivity index across all recorded neurons represents thus a good measure of the action selectivity of the neuronal population as a whole. If this distribution is centered on 0, this indicates that the population has no action selectivity. In contrast, if it is shifted to the right or the left, this indicates that the population fires more or less during action X than during action Y, respectively. Wilcoxon signed-rank tests were used to measure significant shifts from zero in distribution plots. Neuronal activity was normalized by a z-score transformation and population activity was constructed by averaging the z-score activity of all coding neurons. To provide a better visualization of the data, population activity graphs were smoothed with a Gaussian filter.

#### Video analysis of locomotion

Concurrent with *in vivo* neuronal recording, we also measured animals’ position and movement during sampling trials using LEDs mounted on the headstage. These LEDs were detected by an overhead camera coupled to a video tracking system (CinePlex, Plexon Inc, Dallas, TX; 30 frames/s, 1.5 mm spatial resolution). Because locomotion toward the lever could contribute to OFC firing activity, we identified for all sampling trials periods when rats moved in the direction of the lever with a velocity greater than 2 cm/s during at least 0.5 s. We compared OFC neuronal activity during periods of locomotion toward the lever with OFC neuronal activity during preceding baseline periods when rats were not moving (i.e., −12s to-9s before the action) and during periods when rats performed the lever press action itself (i.e.,−1s to 1s after the press) (Figure S2).

### Histology

At the end of the study, histological procedures were used to identify the location of all wire tips used to record neurons. Under anesthesia, an anodal current (50 μA for 5 s)was passed through each microwire to create a small iron deposit. Animals were then perfused with 3.6% paraformaldehyde and 5% potassium ferricyanide solution to stain the iron deposits. The brains were cut into 50 μm coronal sections which were mounted on slides. Only neurons recorded from wires that were within the lateral (LO) and ventral (VO) parts of the OFC were included in data analyses (Figure S1).

### Drugs

Cocaine hydrochloride (Coopération Pharmaceutique Française, Melun, France) was dissolved in 0.9% NaCl, filtered through a syringe filter (0.22 μm) and stored at room temperature. Drug doses were expressed as the weight of the salt. Sodium saccharin (Sigma-Aldrich, France) was dissolved in tap water at room temperature (21 ± 2°C). Sweet solutions were renewed each day.

## AUTHOR CONTRIBUTIONS

Conceptualization and Methodology, K.G. and S.H.A.; Investigation and Formal analysis, K.G.; Writing – Review & Editing, K.G. and S.H.A. wrote the paper.

## ACKNOWLEDGEMENTS

This work was supported by the French Research Council (CNRS), the Université de Bordeaux, the French National Agency (ANR-2010-BLAN-1404-01, ANR-12-SAMA-003 01), the Conseil Régional d’Aquitaine (CRA20101301022; CRA11004375/11004699) and the NRJ Foundation. We thank Ourida Gaucher, Jean-Philippe Fougères, Audrey Durand and Caroline Vouillac-Mendoza for administrative and technical assistance. We also thank Etienne Gontier from the Bordeaux Imaging Center (BIC) for giving us access to his laboratory facility for brain perfusion.

